# HPC-based genome variant calling workflow (HPC-GVCW)

**DOI:** 10.1101/2023.06.25.546420

**Authors:** Yong Zhou, Nagarajan Kathiresan, Zhichao Yu, Luis F. Rivera, Manjula Thimma, Keerthana Manickam, Dmytro Chebotarov, Ramil Mauleon, Kapeel Chougule, Sharon Wei, Tingting Gao, Carl D. Green, Andrea Zuccolo, Doreen Ware, Jianwei Zhang, Kenneth L. McNally, Rod A. Wing

## Abstract

A high-performance computing genome variant calling workflow was designed to run GATK on HPC platforms. This workflow efficiently called an average of 27.3 M, 32.6 M, 168.9 M, and 16.2 M SNPs for rice, sorghum, maize, and soybean, respectively, on the most recently released high-quality reference sequences. Analysis of a rice pan-genome reference panel revealed 2.1 M novel SNPs that have yet to be publicly released.

---

Single-nucleotide polymorphisms (SNPs) are the most common type of genetic variation used for studying genetic diversity among living organisms^1^, and are routinely detected by mapping resequencing data to a reference genome sequence using various software tools^2-4^. The Genome Analysis Toolkit (GATK), one of the most popular software tools developed for SNP identification, has been widely used for SNP detection for many species and can be compiled on multiple computing platforms^5,6^. Although vast amounts of resequencing data have contributed significantly to the study of genetic diversity, at least three challenges remain to be solved for the speed and efficiency of SNP detection. First, the exponential increase in sequencing and resequencing data requires intelligent data management solutions and compressed data formats to reduce storage^7,8^; second, data analysis needs flexible workflows and monitoring tools for high-throughput detection and debugging^9^; and third, modern high-performance computing (HPC) architectures are needed to complete jobs efficiently^10,11^. To address these (and other) bottlenecks, we developed an open source HPC-based automated and flexible genome variant calling workflow for (i.e., HPC-GVCW) for GATK, and tested it on four major crop species (i.e., rice, sorghum, maize, and soybean) using publicly available resequencing data sets and their most up-to-date (near) gap-free reference genome releases.

HPC-GVCW was designed into four phases: (1) mapping, (2) variant calling, (3) call set refinement and consolidation, and (4) variant merging (Supplementary Figure 1a, Supplementary Figure 2, Supplementary Table 1, and Supplementary Note 1). The workflow was also designed to run on various computational platforms, including high-performance computers, hybrid clusters, and high-end workstations (Supplementary Figure 1b). For phase 3, a data parallelization algorithm “Genome Index splitter” (GIS release1.2, https://github.com/IBEXCluster/Genome-Index-splitter) was developed to split chromosome pseudomolecules into multiple chunks (Supplementary Figure 2e and Supplementary Note 1). Leveraging this algorithm ensures that the creation of disjoint variant intervals is optimized based on genome size and computational resources, thereby preventing the underutilization of resources and the reduction of execution times. Of note, each of the four phases is independent of one another, flexible, and automatically scalable across multiple nodes and platforms, which leads to an efficient SNP calling workflow (Supplementary Note 1).

HPC-GVCW is fully open resource, with all scripts, workflows, and instructions available at GitHub (https://github.com/IBEXCluster/Rice-Variant-Calling) (Supplementary Note 1). In addition, to enhance the flexibility of computing platforms and applications, robust containerization solutions, including Docker^12^ and Singularity^13^ were developed (see Data availability).

To evaluate the precision and execution time performance of GVCW, we assessed the workflow across three computational platforms -i.e., supercomputer, clusters, and high-end workstations, using a subset of the 3K-RGP dataset^14^ (n = 30) mapped to the IRGSP RefSeq^15^, and observed a 83-94% identical call rate when compared with previously published results^16^ and across different platforms (Supplementary Figure 3a). Next, we interrogated the workflow on much larger resequencing data sets from multiple crop species data sets (rice [n=3,024]^14^, sorghum [n=400]^17^, maize [n=282]^18^, and soybean [n=198]^19^) on supercomputers and clusters, where we found identical call rates of between 77.8%-83.2% as compared with published results^14,17^, except for maize, where we identified 167.6 M of SNPs, of which 41.9 M overlapped with published results^18^ (Supplementary Figure 3b-d, Supplementary Table 2, and Supplementary Note 2). Execution time performance showed that GVCW can efficiently call SNPs across whole genomes in 5-10 days depending on the number of resequencing samples, genomes size and computational platform (Supplementary Table 3 and Supplementary Note 2). In short, our benchmarking exercise confirms that GVCW can be efficiently used to rapidly identify SNPs using large datasets for major crop species.

Since the majority of publicly available SNP data for major crop species has yet to be updated on the recent wave of ultra-high-quality reference genomes coming online, we applied GVCW to call SNPs, with the same large resequencing datasets, on the most current and publicly available genome releases for rice (i.e., the 16 genome Rice Population Reference Panel)^20-22^, maize (B73 v4 and v5)^23^, sorghum (Tx2783, Tx436, and TX430)^24^, and soybean (Wm82 and JD17)^25^. The results, shown in Table 1, revealed that an average of 27.3 M (rice), 32.6 M (sorghum), 168.9 M (maize), and 16.2 M (soybean) SNPs per genome could be identified for each crop species, of which 3.0 M (rice), 7.8 M (sorghum), 4.4 M (maize), and 6.1 M (soybean) SNPs are located in exons using SNPEff^26^ (Table 1 and Supplementary Table 4). Of note, SNP data for rice (i.e., ARC, N22, AZU, IR64, IRGSP, MH63 ZS97) and sorghum (Tx2783) genome data sets can be visualized at the Gramene (https://oryza.gramene.org/) and Sorghumbase (https://sorghumbase.org/) web portals, respectively (Supplementary Figure 4). In addition, all SNP data produced for this 24-genome reference set have been publicly released through the SNP-Seek (https://snp-seek.irri.org/), Gramene (www.Gramene.org), and KAUST Research Repository (KRR, https://doi.org/10.25781/KAUST-12WKO)^27^ public databases for immediate access (also see Data availability).

**Table 1.** Number of SNPs identified across four major crop species using their most recent public genome releases.

| Species | Reference Genome | Acronyms | GenBank ID | Number of SNPs | SNPs in exons | SNPs in 3' UTR | SNPs in 5' UTR |
| --- | --- | --- | --- | --- | --- | --- | --- |
| Rice<br>(*Oryza sativa*)<br>Genome Size: ~400 Mb | GJ-temp: IRGSP | IRGSP | GCF_001433935.1 | 26,516,112 | 3,060,410 | 319,632 | 232,847 |
|  | GJ-subtrp: CHAO MEO | CM | GCA_009831315.1 | 27,024,845 | 3,069,706 | 356,381 | 233,761 |
|  | GJ-trop1: Azucena | AZ | GCA_009830595.1 | 27,316,403 | 3,081,793 | 345,485 | 226,235 |
|  | GJ-trop2: KETAN NANGKA | KN | GCA_009831275.1 | 27,331,337 | 3,031,741 | 335,086 | 219,831 |
|  | cB: ARC 10497 | ARC | GCA_009831255.1 | 27,286,525 | 2,984,499 | 324,769 | 211,937 |
|  | XI-1A: ZhenShan97RS3 | ZS97 | GCA_001623345.2 | 27,439,649 | 3,504,390 | 573,128 | 406,815 |
|  | XI-1B1: IR 64 | IR64 | GCA_009914875.1 | 27,084,312 | 2,822,657 | 311,142 | 203,724 |
|  | XI-1B2: PR 106 | PR106 | GCA_009831045.1 | 27,461,145 | 3,029,730 | 343,797 | 224,081 |
|  | XI-2A: GOBOL SAIL | GS | GCA_009831025.1 | 27,608,213 | 2,885,485 | 293,846 | 198,221 |
|  | XI-2B: LARHA MUGAD | LM | GCA_009831355.1 | 27,974,114 | 2,921,223 | 307,604 | 206,271 |
|  | XI-3A: LIMA | LIMA | GCA_009829395.1 | 27,053,048 | 2,838,843 | 301,480 | 197,894 |
|  | XI-3B1: KHAO YAI GUANG | KYG | GCA_009831295.1 | 27,378,477 | 2,911,252 | 307,567 | 201,613 |
|  | XI-3B2: LIU XU | LX | GCA_009829375.1 | 27,759,204 | 2,939,867 | 311,835 | 213,624 |
|  | XI-adm: MH63RS3 | MH63 | GCA_001623365.2 | 27,503,492 | 3,509,396 | 603,812 | 422,385 |
|  | cA1: N22 | N22 | GCA_001952365.3 | 27,594,493 | 3,019,972 | 328,996 | 229,046 |
|  | cA2: NATEL BORO | NABO | GCA_009831335.1 | 28,044,207 | 2,979,119 | 312,640 | 212,806 |
| Sorghum<br>(*Sorghum bicolor*)<br>(Genome Size: ~600 Mb) | BT623v3.1 | - | GCF_000003195.3 | 32,698,281 | 1,078,742 | 793,513 | 675,414 |
|  | Tx2783 | - | GCA_903166285.1 | 32,537,001 | 752,298 | 327,512 | 205,336 |
|  | Tx436 | - | GCA_903166325.1 | 32,748,001 | 868,964 | 422,070 | 247,710 |
|  | Tx430 | - | GCA_003482435.1 | 35,102,930 | 1,194,497 | 360,556 | 236,007 |
| Maize<br>(*Zea mays*)<br>Genome Size: ~2000 Mb | B73v4 | - | GCF_000005005.2 | 167,604,407 | 5,789,626 | 3,758,096 | 3,413,940 |
|  | B73V5 | - | GCA_902167145.1 | 170,004,877 | 3,073,808 | 1,325,232 | 1,023,768 |
| Soybean<br>(*Glycine max*)<br>Genome Size: ~1000 Mb | Wm82.a2.v1 | - | Gmax 275 | 15,994,704 | 812,611 | 267,541 | 194,096 |
|  | JD17 | - | GCA_021733175.1 | 16,341,705 | 569,416 | 213,129 | 147,393 |

Having the ability to map large-scale resequencing datasets rapidly (e.g., 3K-RGP) to multiple genomes (e.g., the 16-genome RPRP dataset), GVCW opens the possibility to rigorously interrogate population-level pan-genome datasets for core, dispensable, and private variants (e.g., SNPs, insertions/deletions, inversions).

Our analysis of the 3K-RGP dataset^14^ mapped to the 16-genome RPRP dataset^21^ revealed a core genome of 314.1 Mb, an average dispensable genome of 56.55 Mb, and a private genome of ∼745 Kb/genome (see methods for definitions), that contain ∼22.4 M, 3.2 M and 33.8 K SNPs, respectively (Supplementary table 5, Supplementary Figure 5, and Supplementary Dataset 1-3). We found that an average of 36.5 Mb of genomic sequence is absent in a single rice genome but is present in at least one of the other 15 RPRP data sets, which is equivalent to ∼2.1 M SNPs (Figure 1, Supplementary Figure 5, and Supplementary Table 5). For example, when considering the most widely used reference genome for rice, i.e., in the IRGSP RefSeq^15^, a total of ∼36.6 Mb of genomic sequence is completely absent in the IRGSP RefSeq but is found spread across at least one of the 15 genomes (∼2.43 Mb/genome), and includes ∼2.3 M previously unidentified SNPs (Figure 1, Supplementary Table 5).

**Figure 1.**
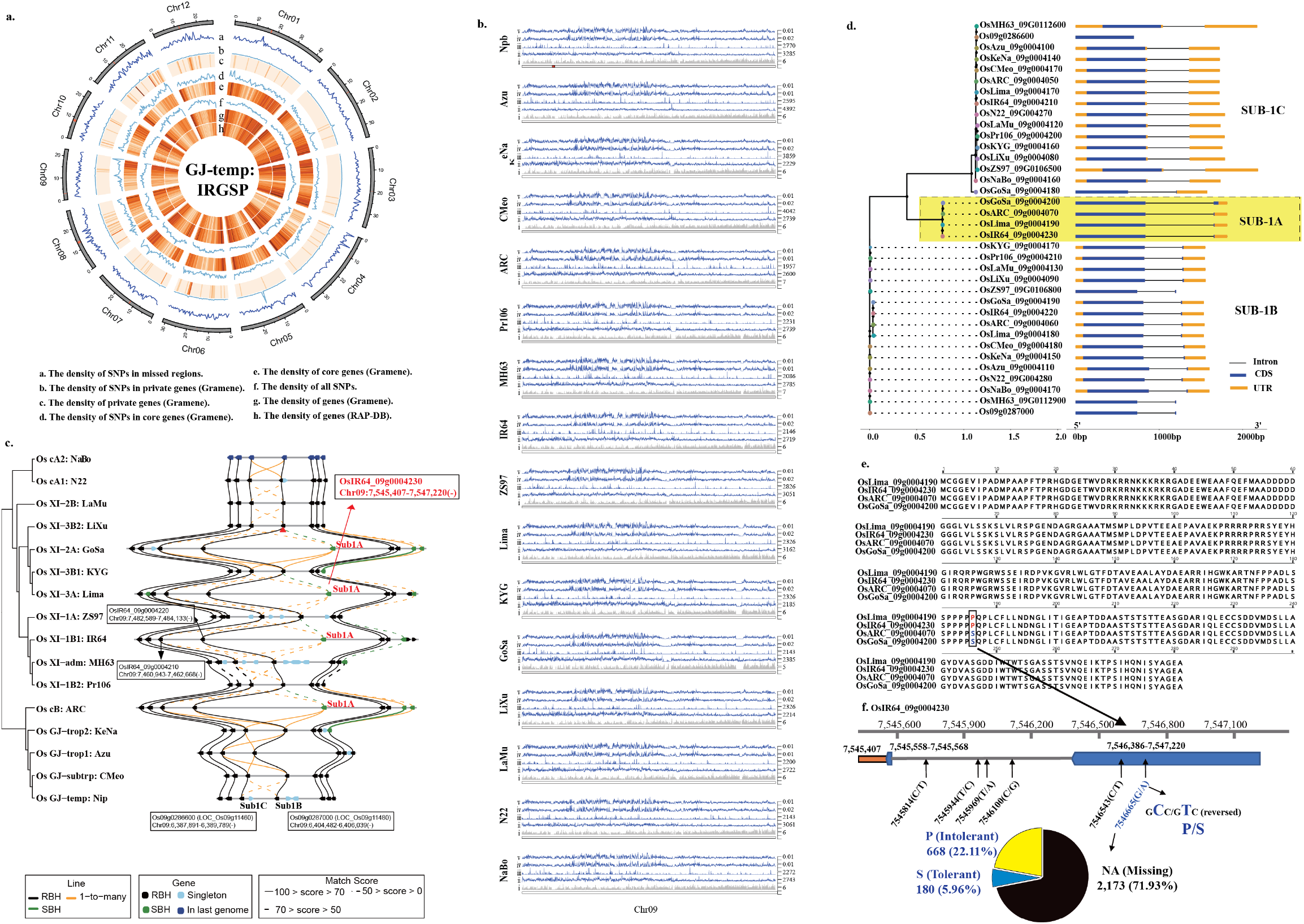
Rice Population Reference Panel (RPRP)^21^ pan-genome variant analysis. **a**, Circos plot depicts the distribution of genomic attributes along the IRGSP RefSeq (window size = 500 Kb). **b**, Comparison of genomic attributes, i.e., genes, SNPs, *Pi*, and *Theta* on chromosome 9 across the 16 RPRP pan-genome data sets (window size = 10 Kb). **c**, Rice Gene Index (RGI) comparison of the *Sub* loci across the 16 RPRP pan-genome data set. **d**, Phylogenetic analysis of *Sub1A, Sub1B*, and *Sub1C* across the 16 RPRP pan-genome data set. **e**, Amino acid alignment of the *Sub1A* gene across the RPRP. **f**, Survey of SNPs within the *Sub1A* gene across the 3K-RGP resequencing data set. This analysis revealed the genomic status of the *Sub1A* gene (presence/absence; submergence tolerance/intolerance) across the 3K-RGP data set.

Performing a similar analysis on gene content using the Rice Gene Index (RGI) database^**22**^ enabled us to identify an average of 24,700, 6,577, and 293 core, dispensable and private homologous gene groups (see methods for definitions) across the 16-genome RPRP data set, respectively (Supplementary Table 5 and Supplementary Dataset 4), equating to 5.5 M SNPs (2.4 M exonic), 0.8 M SNPs (0.2 M exonic), and 37.8 K SNPs (9.6 K exonic) (Figure 1, Supplementary Figure 6 and Supplementary Table 5), respectively. Importantly, on average, a total of ∼10.3 K genes present in 15 of the 16 RPRP genomes (687 genes/genome) are absent in a single RPRP genome, and equates to ∼1.4 M SNPs (0.38 M exonic) (Figure 1, Supplementary Figure 6 and Supplementary Table 5). Again, taking the IRGSP RefSeq as an example, a total of 9,812 genes (accounting for 1.3 M SNPs [0.36 M exonic]) detected in 15/16 of the RPRP genomes are absent in the IRGSP RefSeq (Figure 1 and Supplementary Table 5).

Many of the genes and SNPs identified in our pan-genome variant analysis have yet to be tested for their contributions to agronomic performance and biotic and abiotic stresses. For example, prolonged submergence during floods can cause significant constraints to rice production resulting in millions of dollars of lost farmer income^28^. One solution to flooding survival has been to cross the *Sub1A* gene, first discovered in a tolerant *indica* derivative of the FR13A cultivar (IR40931-26) in 2006^28^, into mega rice varieties such as Swarna, Sambha Mahsuri, and IR64^28,29^. Our analysis of the *Sub1A* locus across the pan-genome of rice showed that this gene could only be observed in 4 out of 16 genomes in the RPRP data set, including IR64 (Figure 1C-D). Since *Sub1A* is absent in the IRGSP RefSeq, the genetic diversity of this locus can only be revealed through the analyses of reference genomes that contain this gene. Thus, we applied the IR64 reference as the base genome for SNP comparisons, and identified a total of 26 SNPs in the *Sub1A* locus across 3K-RGP (Supplementary Dataset 5), 6 of which have minor allele frequencies (MAF) greater than 1% (Figure 1F), including a previously reported SNP (7,546,665-G/A) that resulted in a non-conservative amino acid change from serine (S, *Sub1A-1*, tolerance-specific allele) to proline (P, *Sub1A-2*, intolerance-specific allele)^28^ (Figure 1E-F). The majority of accessions in the 3K-RGP data set (i.e., 2,173) do not contain the *Sub1A* gene, while 848 do, 668 of which (22.11%) have the *Sub1A-2* allele, while 180 accessions (5.96%) contain the *Sub1A-1* allele (Figure 1F). Understanding the genetic diversity of the *Sub-1A* gene at the population level helps us understand and filter variants that are predicted to show flooding tolerance across the 3K-RGP, which could be further applied to precise molecular-assisted selection (MAS) breeding programs. In addition, such pan-genome analyses may also reveal new variants that could provide valuable insights into the molecular mechanisms of flooding tolerance.

With the ability to produce ultra-high-quality reference genomes and population-level resequencing data -at will -accelerated and parallel data processing methods must be developed to efficiently call genetic variation at scale. Some of the accelerated workflows include Sentieon^30^ (a commercial license), Clara Parabricks^31^ (NVIDIA GPU-based infrastructure), Falcon^32^ (hybrid FPGA-CPU cloud-based software), and DRAGEN-GATK^33^ (open source software recently made available through the Broad Institute cloud platform, https://broadinstitute.github.io/warp/) are the examples. These workflows require special hardware (e.g., GPUs, FPGAs) or a cloud computing platform to accelerate the data processing, and/or can be expensive to purchase. To address such limitations, we developed a publicly available high-performance computing pipeline -i.e., the Automated and Flexible Genome Variant Calling Workflow (HPC-GVCW) -for one of the most popular SNP callers – GATK^5,6^. GVCW is supported across diversified computational platforms, i.e., desktops, workstations, clusters, and other high-performance computing architectures, and was containerized for both Docker^12^ and Singularity^13^, for reproducible results without reinstallation and software version incompatibilities.

Comparison of SNP calls on identical data sets (i.e., rice 3K-RGP to the IRGSP RefSeq and 400 samples from Sorghum Association Panel to the BT623v3.1) yielded similar results, however, run times could be reduced from more than six months to less than one week (∼24 times faster), as in the case for rice 3K-RGP^14^. The GVCW pipeline enabled the rapid identification of a large amount of genetic variation across multiple crops, including sorghum, maize, and soybean on the world’s most up-to-date, high-quality reference genomes. These SNPs provide an updated resource of genetic diversity that can be utilized for both crop improvement and basic research, and are freely available through the SNP-Seek (https://snp-seek.irri.org/), Gramene (www.Gramene.org) web portals, and KAUST Research Repository (KRR, https://doi.org/10.25781/KAUST-12WKO)^27^.

Key to our ability to rapidly call SNPs on a variety of computational architectures lies in the design of the HPC environment and the distribution of work across multiple nodes. Our next steps will be to apply GVCW on improved computing platforms, e.g., KAUST Shaheen III with unlimited storage and file number, 5,000 nodes, faster I/O, and tests on larger forthcoming data sets (https://www.kaust.edu.sa/en/news/kaust-selects-hpe-to-build-powerful-supercomputer). In addition to GATK, other SNP detection strategies such as the machine learning based tool “DeepVariant”^2^, which shows better performance in execution times with human data^4^, has yet to be widely used in plants. With a preliminary analysis of rice 3K-RGP dataset, “DeepVariant” identified a larger number of variants at a similar or lower error rate compared to GATK (https://cloud.google.com/blog/products/data-analytics/analyzing-3024-rice-genomes-characterized-by-deepvariant). To test how artificial intelligence (AI) can be used to improve food security by accelerating the genetic improvement of major crop species, we plan to integrate “DeepVariant” into our HPC workflow to discover and explore new uncharacterized variation. In addition, we also plan to apply similar pan-genome strategies on more species beyond rice, sorghum, maize and soybean to discover and characterize hidden SNPs and diversity, which could provide robust and vital resources to facilitate future genetic studies and breeding programs.

## Supporting information

HPC-GVCW.Manuscript.BioRxiv.Supplementary

HPC-GVCW.Manuscript.BioRxiv.Tables

## Supplementary files

Supplementary Note 1: Automated and Flexible computing genome variant calling workflow (HPC-GVCW).

Supplementary Note 2: GVCW workflow performance. Supplementary Note 3: Acronyms used in this manuscript.

## Data availability

All sequence data are available in public databases as follows.

All genome assemblies for rice, sorghum, maize, and soybean were retrieved from NCBI (Table 1), except for Wm82.a2.v1, which is available at the Phytozome (https://phytozome-next.jgi.doe.gov/info/Gmax_Wm82_a2_v1).

Genome resequencing data sets for rice (n=3,024), sorghum (n=400), maize (n=282), and soybean (n=198) were retrieved from NCBI via BioProject accession numbers: PRJEB6180 (https://www.ncbi.nlm.nih.gov/bioproject/PRJEB6180),

PRJEB50066 (https://www.ncbi.nlm.nih.gov/bioproject/PRJEB50066),

PRJNA389800 (https://www.ncbi.nlm.nih.gov/bioproject/PRJNA389800), and PRJDB7281 (https://www.ncbi.nlm.nih.gov/bioproject/PRJDB7786) respectively.

SNP datasets for the RPRP pan-genome are publicly available at:

SNP-Seek (https://snp-seek.irri.org/_download.zul);

Gramene (http://ftp.gramene.org/collaborators/Yong_et_al_variation_dumps/);

KAUST Research Repository (KRR, https://doi.org/10.25781/KAUST-12WKO)^27^.

Realignment data sets of near variant regions (cram file format) of the *O. sativa* 16-genome RPRP data set are available through Amazon Web Services (AWS) 3kricegenome bucket at SNP-Seek (https://snp-seek.irri.org/_download.zul).

SNP datasets for sorghum, soybean, and maize are released at Gramene (http://ftp.gramene.org/collaborators/Yong_et_al_variation_dumps/), and KAUST Research Repository (KRR, https://doi.org/10.25781/KAUST-12WKO)^27^.

SNP datasets for sorghum can be visualized from the Sorghumbase web portal (https://www.sorghumbase.org/).

## Code availability

All code for the Automated and Flexible Workflow for Genome Variant Calling (GVCW) is available on the GitHub page: https://github.com/IBEXCluster/Rice-Variant-Calling.

The Docker and Singularity images are available at https://github.com/IBEXCluster/Rice-Variant-Calling/wiki/Docker and https://github.com/IBEXCluster/Rice-Variant-Calling/tree/main/Singularity, respectively.

## Acknowledgments

This research was supported by King Abdullah University of Science & Technology’s Baseline funding and the University of Arizona’s Bud Antle Endowed Chair for Excellent in Agriculture to R.A.W. The authors acknowledge support from the Shaheen Cray XC40 Supercomputing and Ibex heterogeneous cluster platforms at KAUST Supercomputing Laboratory (KSL). The authors acknowledge data availability and visualization at SNP-Seek (https://snp-seek.irri.org/), and Amazon Web Services (AWS) Open Data as a data repository, Gramene (www.Gramene.org), Sorghumbase (https://www.sorghumbase.org/) portals, and KAUST Research Repository.

## Author Contributions Statement

R.A.W. designed and conceived the research. Y.Z., N.K., Z.Y., and L.F.R. led and operated the project. N.K., Y.Z., L.F.R., M.T., and D.C. designed and tested the HPC pipeline. N.K., L.F.R., and C.G. managed the computing platforms. Y.Z., N.K., K.Ma., M.T., S.W., and K.C, operated the data process of the HPC pipeline and visualization. Y.Z., A.Z., Z.Y., and T.G. identified and validated large structure variations. Z.Y., Y.Z., and J.Z. identified orthologous and studied the specific SNPs. K.C., S.W., and D.W. managed the data transfer and availability at Gramene. R.M., D.C., and K.L.M. managed the data transfer and availability at IRRI (SNP-Seek). Y.Z., N.K., Z.Y., and R.A.W. wrote and edited the paper. All authors read and approved the final manuscript.

