## Supplementary material for "HPC-based genome variant calling workflow (HPC-GVCW)": HPC-GVCW.Manuscript.BioRxiv.Supplementary

### Online methods

#### 1. SNP identification workflow

The SNP identification workflow presented here (i.e., genome variant calling workflow (GVCW)) was developed to provide a freely available and containerized high-performance computational platform to run the Genome Analysis Toolkit (GATK) "best practice" software (https://gatk.broadinstitute.org/hc/en-us/sections/360007226651-Best-Practices-Workflows) for the analysis of large resequencing data sets mapped to multiple reference genomes (see Supplementary Note 1 for a detailed description). Briefly, genome resequencing data from multiple crop species was used for quality control, mapping, SNP calling, and multi-sample joint genotyping. Raw Illumina read data was scanned by Fastqc (0.11.8)[1], and trimmed with Trimmomatic (v0.38)[2] with the following parameters: 'ILLUMINACLIP: TruSeq3-PE-2.fa: 2:30:10 LEADING: 3 TRAILING: 3 SLIDINGWINDOW: 4:15 MINLEN: 36'. Trimmed reads were then aligned to their respective high-quality reference genome sequences using BWA-MEM (v0.7.17)[3] under default parameters. Mapped reads with quality scores ≥ 30 were then sorted using SAMTools[4] (v1.8). Duplicate reads were marked and re-grouped using GATK's (v4.1.6)[5] 'MarkDuplicates' and 'AddOrReplaceReadGroups' functions. SNPs for each accession (gVCF) were called using the GATK's HaplotypeCaller[6]. GATK functions 'CombineGVCFs' and 'GenotypeGVCFs' were then used for joint genotyping to produce merged VCFs from gVCFs for each sample by intervals. Finally, SNPs were extracted from the joint genotypes using GATK's 'SelectVariants' and 'VariantFiltration' functions with the following parameters: 'QUAL < 30.0 || QD < 2.0 || MQ < 20.0 || MQRankSum < -3.0 || ReadPosRanKSum < -3.0 || DP < 5.0' to filter for high-quality of SNPs.

#### 2. Sequence data

Twenty-four reference genome sequences, including the latest gap or near gap-free assemblies of rice, sorghum, maize, and soybean, are listed in Table 1. All resequencing data was downloaded from the following public databases: rice 3K-RGP data set (3,024 samples)[7]; sorghum association panel (SAP, 400 samples)[8], maize association mapping panel (AMP, 282 samples)[9], soybean mini-core collection (MCC, 198 samples)[10] (Supplementary Table 3 and Data Availability).

3. SNP annotation

SNPs located in coding regions across the 24 reference genome data sets were identified using their respective annotation files, and functional SNPs were predicted using SnpEff (v5.0e)[11].

4. SNP visualization for rice and sorghum

SNP data for rice (i.e., ARC, N22, AZU, IR64, IRGSP, MH63 ZS97) and sorghum (Tx2783) genome data sets can be visualized at the following web portals, respectively:

Rice: https://oryza.gramene.org/ (Gramene release 6, https://oryza.gramene.org/News)

Sorghum: https://sorghumbase.org/ (Sorghumbase Release 6, https://www.sorghumbase.org/relnotes)

Two examples of putative SNPs that result in premature stop codons are shown in Supplementary Figure 4.

5. Structural variation (SV) update across the 16-genome Rice Population Reference Panel (RPRP)

In 2020, we published an index of large structure variations (>50 bp, SVs) across the 16-genome RPRP[12] that included the MH63RS2 and ZS97RS2 genome assemblies. Here, we updated this index (Supplementary Dataset 2) using the latest gap-free genome assemblies for these genomes - i.e., MH63RS3 and ZS97RS3[13] - using the same methods as previously described[12]. To validate this updated SV index, we randomly selected 50 insertions and 50 deletions across the 16 rice genome RPRP, using the IRGSP RefSeq as the reference and the remaining 15 rice genomes as queries, which included a total of 1,500 entries ($(50 + 50) \times 15 = 1{,}500$).

We then manually validated each SV with alignment information in IGV using raw reads and alignment blocks with Nucmer[14]. SVs were considered valid if the two methods could identify the identical insertion or deletion and resulted in 94.6% of the insertions and 99.3% of the deletions being validated as true SVs (Supplementary Dataset 3).

6. Homologous gene identification across the 16-genome Rice Population Reference Panel (RPRP) based on sequence alignment and syntenic position.

As with SVs above, we also updated our rice gene index (RGI) using updated MH63RS3 and ZS97RS3 gene annotations with the identical pipeline[15]. Briefly, homologous gene sets across the 16-genome RPRP were identified using GeneTribe software[16], by combining protein sequence similarity and collinearity (i.e., synteny) information. Homologous relationships included “reciprocal best hits” (RBHs), “single-side best hits” (SBHs), one-to-many, and singletons. Based on the one-to-one relationships (both RBH and SBH), and considering the collinearity blocks, we removed redundant homologous gene groups to obtain 79,111 non-redundant homologous gene groups. Finally, these non-redundant homologous gene groups were clustered with the “Connected Graph Algorithm”[17] to obtain 41,137 homologous gene groups (Supplementary Dataset 4).

7. Rice pan-genome SNPs analysis

Using the updated SV and RGI data sets in combination with the 16-genome RPRP SNP data set, we conducted a pan-genome SNP analysis to classify genomic regions into core, dispensable, genome-specific, and genome-absent regions[18]. Core regions are defined as sequences that are present in all 16 RPRP genomes. Dispensable regions are defined as sequences that are observed in 2 to 15 of the 16 RPRP genomes. Genome-specific regions are defined as sequences that are present in only one of the 16 RPRP genomes, but absent in the remaining 15. Genome-absent regions are defined as sequences that are not present in one of the 16 RPRP genomes, but are present in at least one of the other 15 genomes. For the presence and absence of genes, we classified homologous gene groups as core, dispensable, specific, and absent genes, representing the same logic flow as large SVs. Bedtools (v2.30.0)[19] subcommand “subtract” was used for core region identification, and the subcommand “intersect” was used for SNP extraction.

### Supplementary Note 1: Automated and flexible high-performance computing based genome variant calling workflow (HPC-GVCW)

1. Background

Next-generation sequencing (NGS) technologies produce massive volumes of data, requiring sophisticated bioinformatics tools to accurately and rapidly identify genetic variation for genetic diversity, population genetics, genome-wide association studies, etc[20,21]. These analyses require computationally demanding operations that are built with multi-stage workflows for various application tools[7] that can be executed on high-performance computing (HPC) platforms[22,23]. For example, many popular bioinformatics applications can only be executed on single nodes due to the tendency to use multi-threaded programming applications[24]. Thus, methods are needed to use parallel data processing protocols to improve workflow execution times, such as those found on HPC platforms[25]. Furthermore, automating such workflows using schedulers can provide robustness and ease of operation[26]. Finally, once such HPC applications are developed, they can be containerized to be easily installed and executed on a variety of HPC computing ecosystems[27].

In practical terms, when implementing a flexible workflow on different systems, many characteristics need to be addressed in different ways[28]. For example, compute-intensive tools may get accelerated through high-frequency processors; memory-intensive applications require large memory nodes; and, parallel file systems, or local storage on high-end workstations, may help in cases of I/O-intensive calculations[29]. Such computational heterogeneity is extremely difficult to implement on single stage of workflows, hence our motivation and need to develop a multi-stage, automated, flexible, and portable workflow for diversified computing platforms.

The Genome Analysis Toolkit (GATK)[5,6] offers a variety of tools and best practice workflows for both per-sample and joint genotyping (a full list of functionalities is available online: https://gatk.broadinstitute.org/hc/en-us/sections/360008481611-4-1-6-0). The per-sample method of genotyping is straightforward, where each sample can be independently processed across a cluster of nodes to reduce run times. However, joint genotyping is more complex and requires both (i) "distributed parallelization," where users can split chromosomes into disjoint intervals called "chunks"[30] that can be independently executed across clusters of nodes using a job scheduler; and (ii) "shared memory parallelization," where GATK manages multiple instances of output collection from individual chunks, and merging them into the correct reference-based order[31,32]. In the first approach, users require algorithms for data parallelization. In contrast, the second approach is limited to system architectures and is predominantly suitable for a shared memory paradigm. In addition, the performance of shared memory parallelization is poor, and thus, a distributed parallelization method[32] is often employed to improve performance.

Consolidation of gVCFs from multiple samples is called "joint calling," and the distributed parallelization approach is used to minimize execution times. However, this approach is challenging for the following reasons: (i) GATK's "CombineGVCFs" is not scalable across node clusters due to application limitations[33], but luckily, GATK does accept one or more genomic intervals, and multiple instances of GATK with disjoint intervals can be executed across multiple nodes (https://gatk.broadinstitute.org/hc/en-us/articles/360035531852-Intervals-and-interval-lists); (ii) the selection of genomic intervals can be challenging because of imbalanced resource utilization due to suboptimal genomic interval size (i.e., when small genomic intervals are completed, GATK waits for the larger genomic intervals to be completed before all partial results can be merged)[34]; and (iii) job management without a scheduler can be a complex problem, especially when the number of genomic intervals increases exponentially[35].

To alleviate many of these issues, a strategy to split whole genome assemblies into chromosomes to run "joint caller" was devised for fonio millet[36] (i.e., "SNPcaller" workflow: https://github.com/IBEXCluster/IBEX-SNPcaller), where overall execution times were significantly reduced, as compared with a shared memory parallelization method. Alternatively, "bcbio-nextgen" uses a joint caller algorithm that can be executed across multiple instances of GATK. However, execution times vary from smaller to larger chromosomes[36], and have not been optimized to split genomes based on chromosome assignment. Therefore, the selection of optimal variant interval sizes has remained unaddressed for widely used algorithms, including both GATK and GenomicsDB[37-40].

Lastly, workflow automation is also highly recommended, especially when: (i) the number of samples is relatively large (*e.g.,* 3,024 resequenced samples); (ii) job dependencies are very complex when dealing with large sample sizes; and (iii) multi-sample joint genotyping has a large number of variant intervals. To simplify these automation and job management challenges, Message Passing Interface (MPI)[41] wrappers can be used. Disjoint datasets (i.e., multiple samples or the computed variant intervals) are scattered across a cluster of nodes mapped via MPI ranks. Thus, concurrent instances of GATK with disjoint datasets can be executed in parallel, and can be managed as a single job with MPI, thereby simplifying job management and automation.

### 2. Automated Genome Variant Calling Workflow (GVCW) Design

The genome variant calling workflow used in this study was designed and automated for large-scale genomic data sets (e.g., 3,024 Rice Genomes Project (3K-RGP)[7]), to significantly reduce manual data management processes such as tracking multiple jobs and dependencies on high-performance computing platforms (HPC). Multiple bioinformatics tools were employed at various workflow phases (also called "job steps"), and the output of earlier job steps can become the input for the next job step, and so forth. In addition, workflow resource requirements, like the number of CPUs, memory size, use of temporary files, and Java heap/stack size, were considered during job scheduling and automation[42,43].

Our variant calling workflow was divided into 4 phases:

**Phase 1: Data pre-processing**

Phase 1 was designed to map clean resequencing reads to a specified reference genome using the BWA-MEM (v0.7.17)[3] aligner. After alignment, the reads are filtered based on quality scores (≥30), and sorted using SAMTools[4]. Next, GATK is used to carry out the realignment of bam files, where all mate-pair information from different sequence reads (*e.g.*, from similar rice genome samples) are synchronized between each read and its mate pair using GATK's "FixMateInformation." Lastly, read duplicates are identified and condensed into a single new read-group using GATK's "MarkDuplicates" and "AddOrReplaceReadGroups" functions. A summary of the data pre-processing workflow is described in Supplementary Figure 2a.

**Phase 2: Variant discovery**

Phase 2 was designed to call variants for each sample and generate gVCFs files. This phase comprises two major steps: first, multiple sorted input files are merged into a single BAM file and (re)sorted into a merged BAM file using SAMTools. Next, SNPs are called simultaneously via a local *de novo*-assembly of haplotypes in an active region using GATK's

"HaplotypeCaller," where a single gVCF file will be generated per sample. The Phase 2 workflow is described in Supplementary Figure 2b.

**Phase3: Call set refinement**

Phase 3 was designed to merge all variants per sample (stored in a gVCF file from Phase 2) into non-redundant joint genotype files (stored as VCF files) by chunks in two steps. First, all gVCF files are merged using the GATK's "CombineGVCFs" function for each chunk. Next, multi-sample joint genotyping is performed based on the merged VCF file for each chunk using the GATK's "GenotypeGVCFs" function.

Both GATK's "CombineGVCFs" and "GenomicsDB" functions are recommended for sample merging. "GenomicDB" is limited in the number of samples can import at one time, and requires samples to be grouped into multiple batches to address this limitation (https://gatk.broadinstitute.org/hc/en-us/articles/360056138571-GenomicsDBImport-usage-and-performance-guidelines). However, "CombineGVCFs" can combine multiple intervals all at one time, once without the need to build a data store (https://gatk.broadinstitute.org/hc/en-us/articles/360035891051-GenomicsDB) as recently demonstrated for 10,588 human samples successfully[44]. Hence, we used "CombineGVCFs" for data parallelization in our workflow.

GATK's "CombineGVCFs" and "GenotypeGVCFs" functions were initially designed to be executed on a single core because of programming limitations[43]. Unfortunately, assembling genotypes across a large number of samples into a single file can take an extremely long time, and requires huge amounts of memory, especially for species with large genome sizes. To address this limitation, the latest version of GATK offers the variant intervals feature (also referred to as "chunks") for both "CombineGVCFs" and "GenotypeGVCFs." To improve the performance in our workflow, disjoint variant intervals (chunks) are used to run in parallel across a node cluster on HPC platforms. In this design, merging variants from multiple samples ("CombineGVCFs") and joint genotyping ( "GenotypeGVCFs") can proceed for each chunk simultaneously.

To optimize the size and number of chunks, an algorithm called "Genome Index Splitter" (GIS) was developed (https://github.com/IBEXCluster/Genome-Index-splitter). This algorithm creates a "chromosome split table" (CST) to index disjoint variant intervals, which can be fine-tuned based on reference genome size and available CPUs (Supplementary Figure 2c-d). Optimal chunks are calculated based on three steps: (1) locate the largest chromosome length in a given reference genome; (2) calculate the fairness of a divisible integer for a given maximum number of cores; and (3) whole genome reference sequences are divided by the optimal integer number, as illustrated in Supplementary Figure 2e.

For example, the CST with the entries as follows: < ChrName, Chunk_no, Start, End >

Chr01 1 1 2277417
Chr01 2 2277418 4554834
Chr01 3 4554835 6832251
Chr01 4 6832252 9109668
Chr01 5 9109669 11387085
Chr02 1 1 2277417
Chr02 2 2277418 4554834
Chr02 3 4554835 6832251
Chr02 4 6832252 9109668
Chr03 1 1 2277417
Chr03 2 2277418 4554834

Once the chunk size is optimized, jobs (both GATK's "CombineGVCFs" and "GenotypeGVCFs" functions) will be distributed and parallelized by chunks (Supplementary Figure 2f-g). GIS has been shown to work across various platforms and different workflows for major crop species over a broad range of genome sizes (GS) - e.g., rice [GS=400Mb] (https://github.com/IBEXCluster/Rice-Variant-Calling/), and wheat [GS=15Gb] (https://github.com/IBEXCluster/Wheat-SNPCaller).
In our recent GIS release (https://github.com/IBEXCluster/Genome-Index-splitter, release 1.2), GIS was updated to use MPI-based data distribution that was also applied successfully for sorghum, maize, and soybean.

**Phase4: Variant matrix**

Phase 4 was designed to generate a genome-wide joint genotype by assembling all disjoint variant intervals from Phase 3 by combining all chunks into a single file using GATK's

"GatherVcfs" option. Chromosome-based SNPs can then be converted into a variant table with "HapMap" format[45] for post-processing analyses (Supplementary Figure 2h).

**Workflow flexibility**

Of note, all software environments and workflow scripts have been "containerized," so the various workflow phases can be effortlessly switched from one system architecture to another (see code availability). For example, data preprocessing of many samples concurrently can be executed in a cluster computing environment (i.e., Phase 1 & 2). The user can then move the results from the cluster to an HPC system for Phase 3 - i.e., "Call set refinement" - without any changes in the existing workflow scripts. This flexibility offers more opportunities to collaborate and utilizes the various computing resources at different organizations. Further, users may migrate workflow environments into any computing platform, including a laptop, cluster computing, and supercomputers, because the software workflow environments are seamlessly supported for any target platform, as illustrated in Supplementary Figure 1.

**Supplementary Note 2: GVCWworkflow performance**

To evaluate the performance of GVCW, we tested the workflow across three computational platforms - i.e., supercomputer, hybrid cluster server, and high-end workstation - and compared parallel SNP call results from a random selection of 30 resequenced genomes mapped to the IRGSP-RefSeq derived from the 3K-RGP[7]. From this assessment, we observed a 83-94% identical call rate across different platforms compared to previously published results[46] (Supplementary Figure 3a), thereby confirming the accuracy of GVCW.

To evaluate execution times of GVCW on larger datasets, we called SNPs on the reference genomes for four major crops (i.e., rice [IRGSP-1.0][47], sorghum [BTx623][48], maize [B73v4][49], and soybean [Gmax275v2.0][50]) using publicly available resequencing data sets from: rice (3K-RGP, 3,024 samples)[7]; sorghum (sorghum association panel [SAP], 400 samples)[8]; maize (maize association panel [MAP], 282 samples)[9]; and soybean (soybean mini-core collection [SMCC], 198 samples)[10]. Using KAUST's Shaheen 2 supercomputer with 30K cores, processing 3,024 resequenced samples (3K-RGP) mapped to a single rice reference genome took 94 hours (i.e., 3.91 days) (Supplementary Table 2). For benchmarking on a hybrid cluster the workflow could be completed for this same data set in 921 hours (i.e., 38 days), and theoretically 55,202 hours (i.e., 6.3 years) on a high-end workstation (Supplementary Table 2). The execution time (CPU hours) on a HPC system with 30,000 cores was 587 times faster compared to 50 cores on a high-end desktop computer.

For the sorghum, maize, and soybean data sets, due to the small number of samples, we only benchmarked GVCW on a hybrid cluster with 3000 cores and found that even for a 2 Gb maize genome[51], that SNP calling for 282 samples could be completed within ten days (Supplementary Table 2).

To assess the accuracy of the SNP calls across the four crop genomes, we compared our results with previously identified SNPs using identical reference genomes. Our benchmarking test identified 26.5 M, 32.7 M, 167.6 M, and 15.9 M SNPs for rice (IRGSP-1.0), sorghum (BTx623), maize (B73 v4), and soybean (Gmax 275 v2.0), respectively (Table 1). Compared to the public SNP data sets available using the same reference genomes for rice 3K-RGP[46] and sorghum 400 SAP[50], we found that 22.8 M (86.3%) and 29.2 M (89.3%) identified SNPs in this study were identical with previous results (Supplementary Figure 3c-d and Supplementary Table 3), which indicates that GVCW yields similar results to previous studies.

**Supplementary Note 3:**

Acronyms used in this manuscript.

| No | Acronyms | Full Name |
|---|---|---|
| 1 | 3K-RGP | The 3000 Rice Genome Project (3,024 samples) |
| 2 | GVCW | Genome Variant Calling Workflow |
| 3 | BAM | Binary Alignment Map |
| 4 | BWA | Burrows-Wheeler Alignment |
| 5 | ChrName | Chromosome Name |
| 6 | Chunk_no | Chunk number, unique integer per chromosome |
| 7 | CPU | Central Processing Unit |
| 8 | End | Chromosome end position |
| 9 | GATK | Genome Analysis ToolKit |
| 10 | gVCF | Genomic Variant Call Format |
| 11 | HPC | High-Performance Computing |
| 12 | I/O | Input/Output |
| 13 | IGV | Integrative Genomics Viewer |
| 14 | INDELs | INsertion–DELetion mutations |

| | | |
|---|---|---|
| 15 | IRGSP | The International Rice Genome Sequencing Project |
| 16 | KAUST | King Abdullah University of Science and Technology |
| 17 | MPI | Message Passing Interface |
| 18 | NGS | Next Generation Sequencing |
| 19 | RBHs | Reciprocal Best HitS |
| 20 | RefSeq | Reference Sequence |
| 21 | RPRP | Rice Population Reference Panel |
| 22 | SAM | Sequence Alignment/Map |
| 23 | SAP | sorghum association panel |
| 24 | SBHs | Single-side Best HitS |
| 25 | SNP | Single Nucleotide Polymorphism |
| 26 | Start | Chromosome starting position |
| 27 | SVs | Structural Variants |
| 28 | T2T | Telomere-to-Telomere |
| 29 | VCF | Variant Call Format |

### Supplementary Tables

Supplementary Table 1. Summary of workflow phases.

Supplementary Table 2. Results of variant detection based on the automated workflow for rice, sorghum, maize, and soybean.

Supplementary Table 3. Performance of the genome variant calling workflow (w/GATK4) for rice, sorghum, maize and soybean.

Supplementary Table 4. SNPEff annotations for rice, sorghum, maize, and soybean.

Supplementary Table 5. The number of SNPs was identified by using rice RPRP references.

### Supplementary Figures

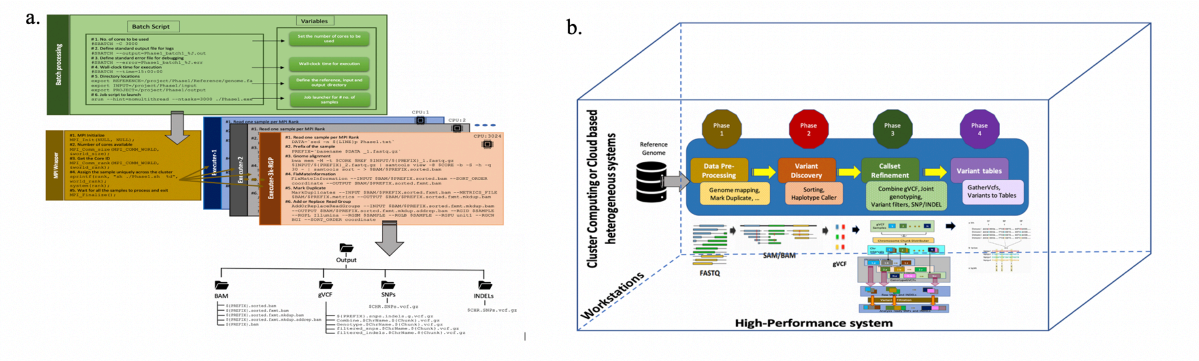


Supplementary Figure 1: Automated and flexible genome variant calling workflow (GVCW) design for a. HPC systems, and b. diversified system architectures.

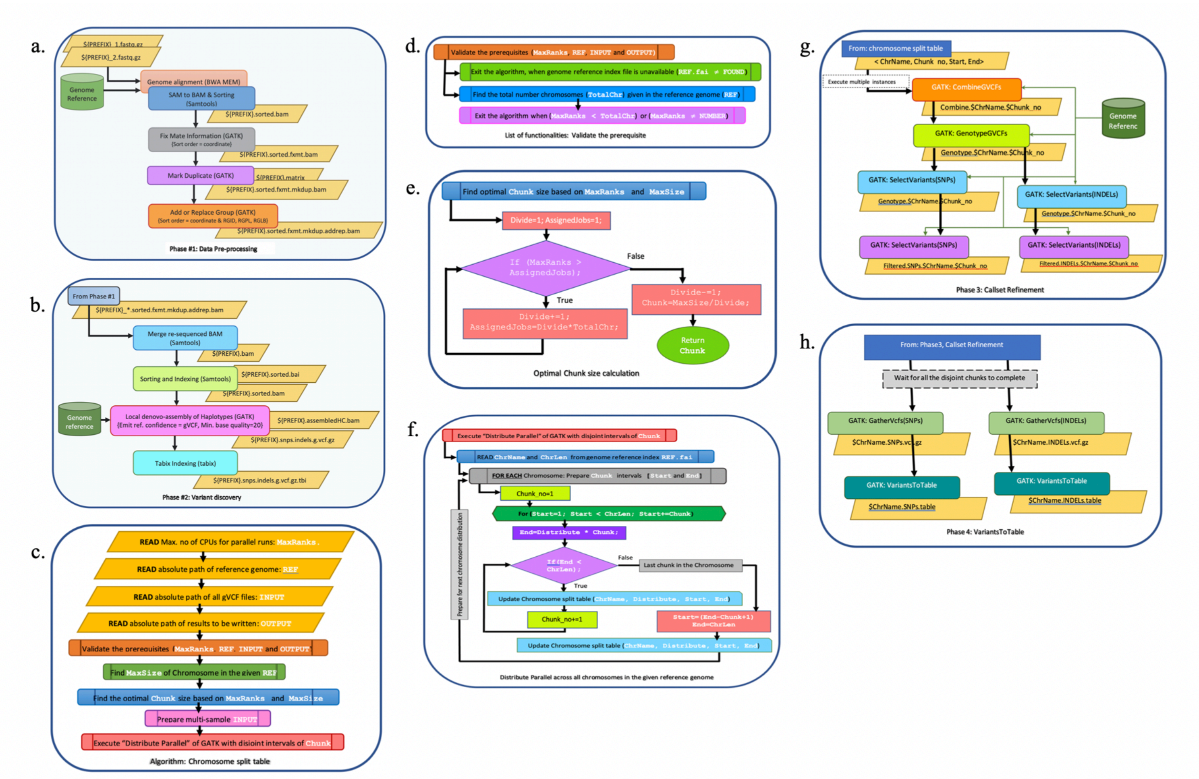


Supplementary Figure 2. GVCW data processing. a. Data pre-processing; b. Variant discovery; c. Chromosome split table for call set refinement; d. Conditions and prerequisites for chromosome split table creation; e. Optimal chunk size calculation; f. Parallel distribution across all the chromosomes for a given reference genome; g. Call set refinement; h. Variants to the table.

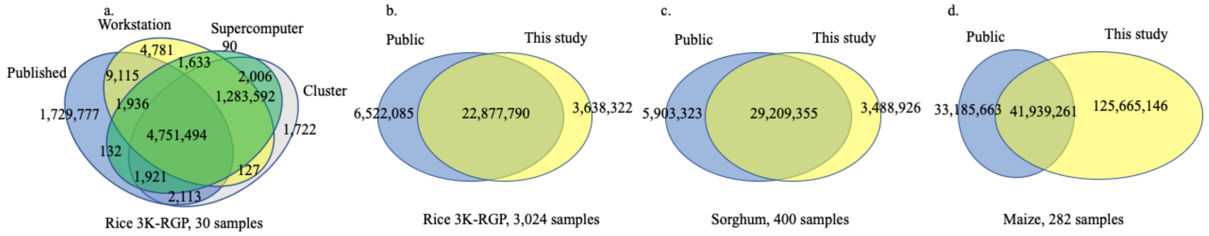


Supplementary Figure 3. Venn diagrams show comparisons of SNP calls for different datasets, i.e., a. Rice (n=30); b. 3K-RGP full datasets for rice (n=3,024); c. Sorghum (n=400); d. Maize (n=282).

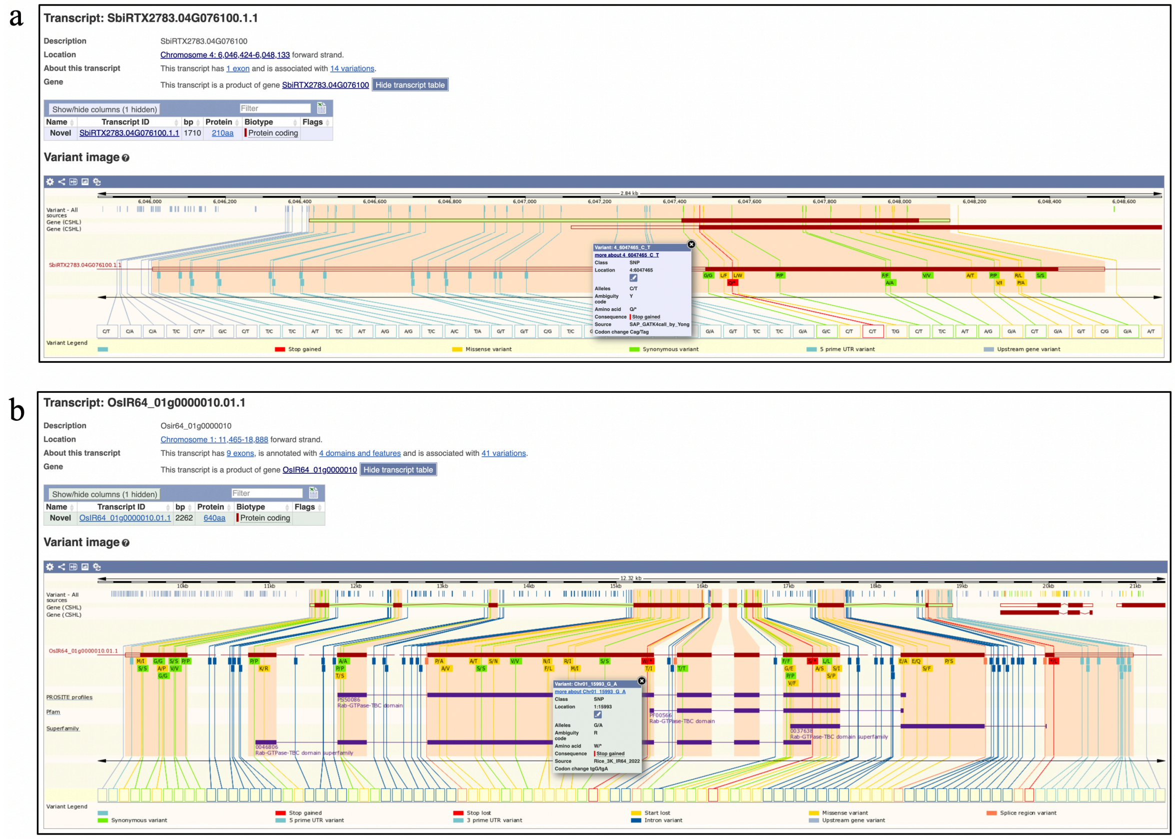


Supplementary Figure 4. SNP visualization of two putative SNPs that resulted in premature stop codons in sorghum (Tx2783) and rice (IR64). a. One C→T transition (Chr04, 6,047,465) for gene SbiRTX2783.04G076100 in the T2783 sorghum genome. b. One G→A transition (Chr01, 15,993) for gene OsIR64_010000010 in the IR64 rice genome.

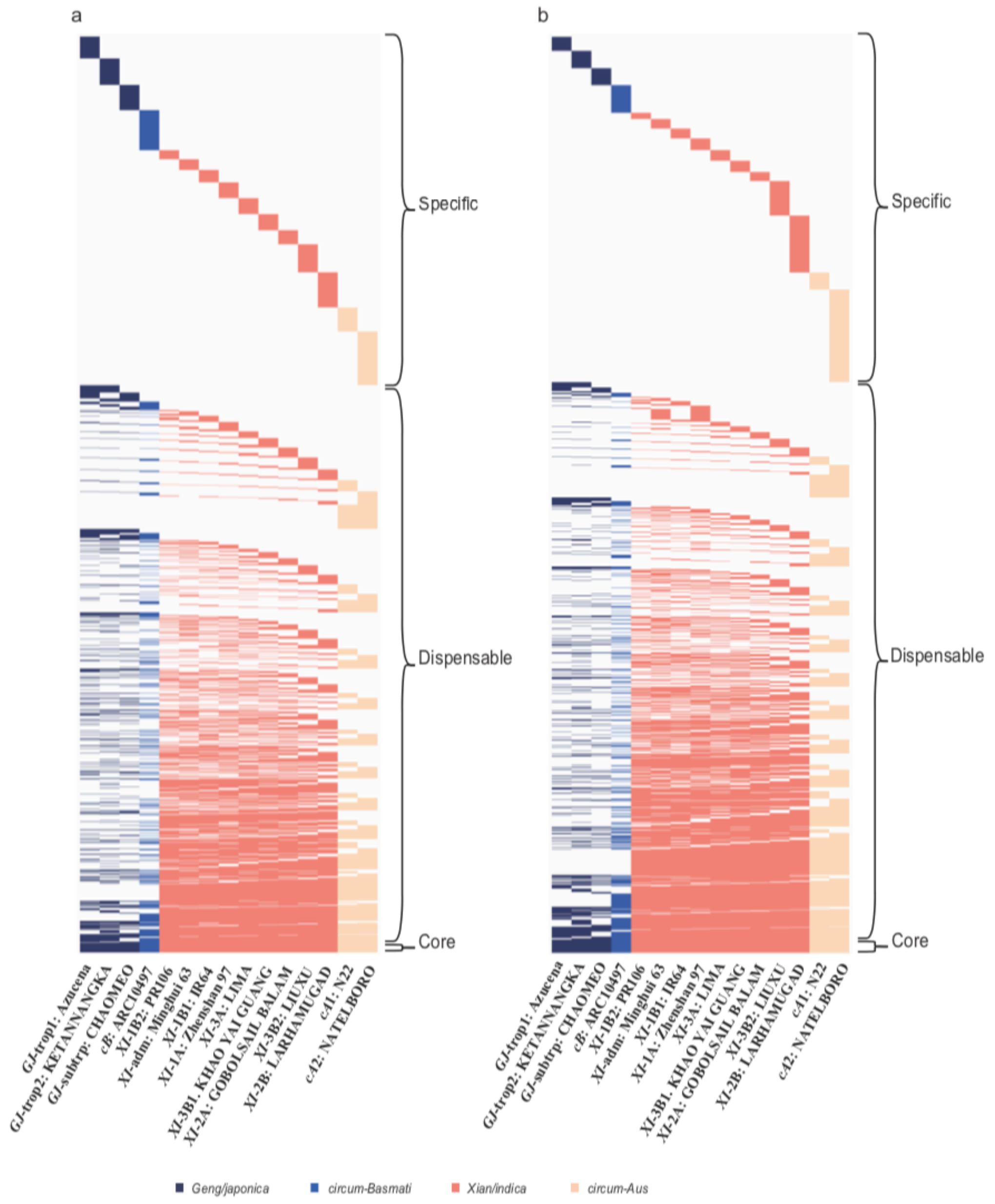


Supplementary Figure 4. Large structure variation (> 50 bp) analysis of pan-genomes Rice Population Reference Panel (RPRP).
a. The insertions and b. deletions were shown, respectively.

Supplementary Figure 5. Large structural variation (> 50 bp) analysis of the 16-genome Rice Population Reference Panel (RPRP). a. Insertions, b. Deletions.

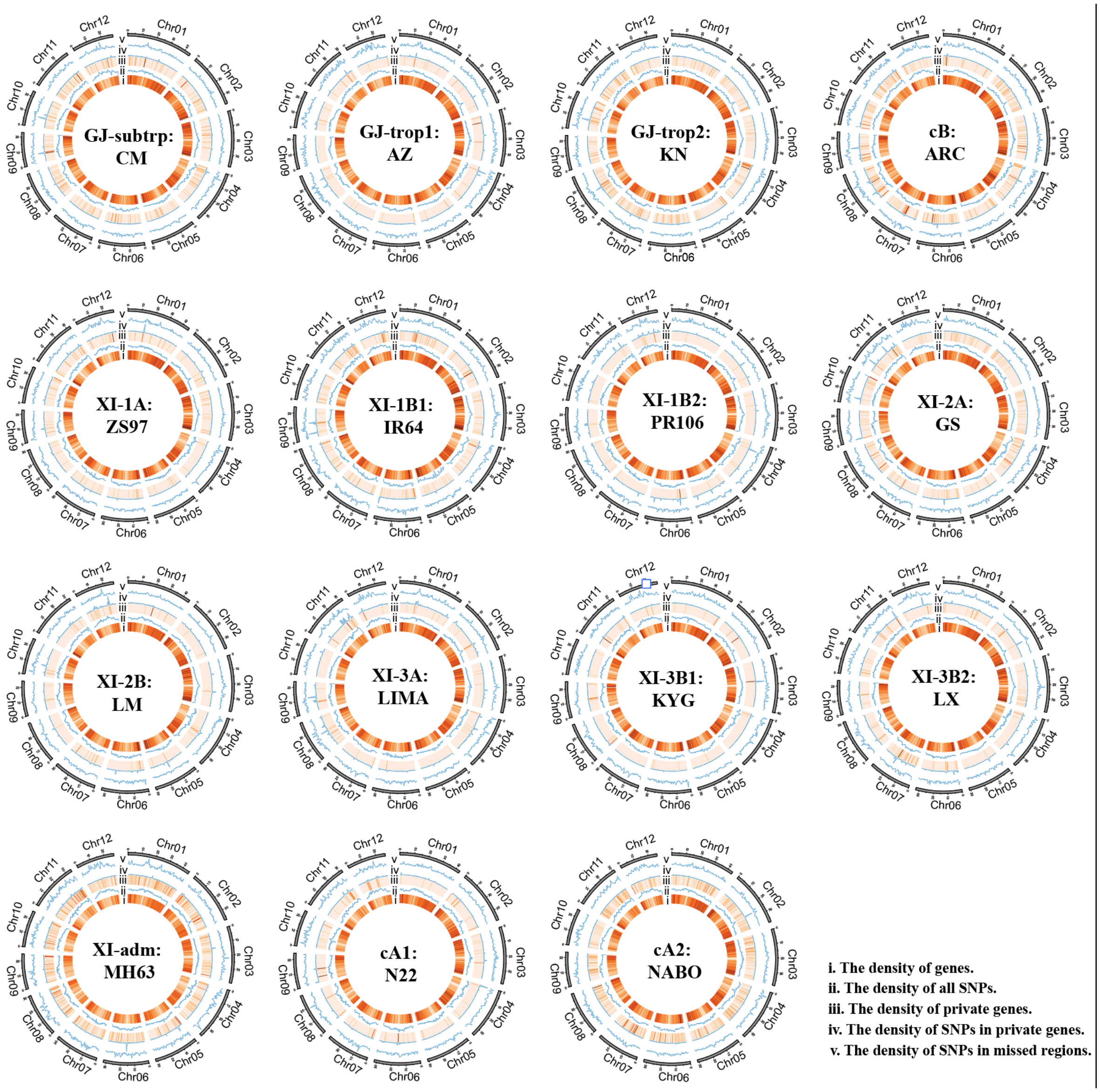


Supplementary Figure 6. Circos plots depict the distribution of genomic attributes along the 12 chromosomes of the 16-genome RPRP data set (window size = 500 Kb).
